# Structures of vesicular stomatitis virus glycoprotein G alone and in complex with a neutralizing antibody

**DOI:** 10.1101/2025.01.13.632142

**Authors:** Marie Minoves, Malika Ouldali, Laura Belot, Stéphane Roche, Lefteris Zakardas, Guy Schoehn, Yves Gaudin, Aurélie Albertini

## Abstract

VSV G mediates viral entry *via* endocytosis. In the endosome, G undergoes a pH-dependent conformational change from pre- to post-fusion state, catalyzing membrane fusion. So far, no complete structure of G has been reported. We report cryo-EM structures of G, isolated from virions using detergent, alone and in complex with neutralizing antibody FAb that binds G in all conformations. The post-fusion structure reveals novel details about the organization of the C-terminal part of the ectodomain, showing that it undergoes conformational rearrangement and stabilizes the post-fusion trimer by nesting into a groove between adjacent fusion domains. The fusion loops are visible inside the micelle, which is not the case of the transmembrane domains, suggesting that they are rather mobile. Structures of G-FAb complex show that the epitope belongs to a conserved antigenic site. This work has potential implications for vaccine development and oncolytic virotherapy.

## Introduction

Vesicular stomatitis virus (VSV) is the prototype of the Vesiculovirus genus in the Rhabdoviridae family. It is an enveloped, bullet-shaped virus capable of infecting a wide range of hosts, including mammals and insects, and is responsible for epizootic outbreaks in cattle in North America ^1^. Its negative-strand RNA genome encodes five structural proteins among which the glycoprotein (VSV G) is the sole transmembrane protein. VSV G plays a crucial role in initiating the viral cycle. It is responsible for both receptor recognition and membrane fusion. Interaction of VSV G with a cellular receptor, particularly members of the LDL-R family ^2,3^, triggers clathrin-mediated endocytosis of VSV ^4,5^. Then, within the acidic environment of the endosome, VSV G catalyzes fusion between viral and endosomal membranes. Indeed, VSV G undergoes a low pH-induced fusogenic conformational change from a pre- toward a post-fusion state. This structural transition results in the transient exposure of hydrophobic motifs (called fusion loops) capable of interacting with the target membrane and destabilizing it, ultimately leading to its fusion with the viral envelope. A particularity of VSV G, shared with other rhabdovirus glycoproteins, is that there is a pH-dependent equilibrium between the pre- fusion state ^6–8^, the post-fusion state, and intermediate conformations on the conformational change pathway. Consequently, the structural transition induced by low pH is reversible^9–11^, which is not the case for other viral fusogens whose pre-fusion conformation is metastable ^12,13^.

VSV G is a type I transmembrane protein. After cleavage of the amino terminal signal peptide, the mature glycoprotein is 495 amino acids (aa) long, the major part being its 446 residues long ectodomain (G_ect_). To date, the three-dimensional (3D) structures of VSV G soluble ectodomains have been determined using X-ray crystallography in both the pre-fusion ^14,15^ and post-fusion states ^16^. These soluble ectodomains were obtained either by proteolytic cleavage at the viral surface or through recombinant expression, with the membrane anchor removed in all cases ^8^. Crystal structures of VSV G_ect_ are available in both pre-fusion ^14,15^ and post-fusion trimeric conformations ^16^. These structures revealed that VSV G shares a common fold with other viral fusion glycoproteins (including gB of herpesviruse ^17^ and gp64 of baculoviruses ^18^), which defines class III membrane fusion proteins ^19,20^. Subsequently, crystal structures of Chandipura virus glycoprotein (CHAV G, another glycoprotein of the Vesiculovirus genus) soluble ectodomain were also solved, corresponding to the post-fusion trimeric conformation ^21^ and two monomeric intermediates ^22^. More recently, the structure of VSV G_ect_ in complex with its cellular receptor binding domains (*i.e.* CR2 and CR3 cysteine-rich domains of the low-density lipoprotein receptor (LDL-R)) was resolved, revealing the organization of the receptor-binding site on VSV G ^2^.

G_ect_ includes three distinct structural domains (nomenclature defined in **Fig. 1a**): the fusion domain (FD), the trimerization domain (TrD), and the pleckstrin homology domain (PHD) that keep their structure during the conformational change. These domains are connected by hinge regions (named R1, R2, R3, R4 and R5) that refold during the structural transition, and therefore reposition structural domains ^23,24^. Finally, the C-terminal domain (CTD) connects the ectodomain to the transmembrane domain (TMD) (**Fig. 1a**).

**Fig. 1.**
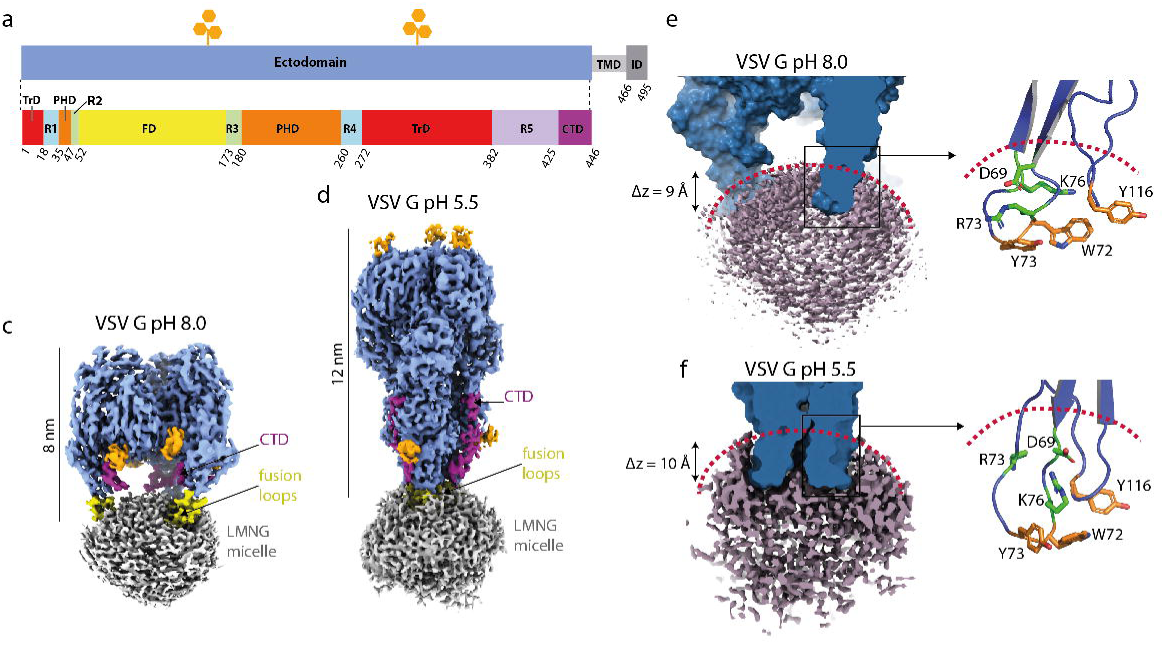
VSV G cryo-EM structures in its pre- and post-conformations. **a** Schematic diagram of VSV G sequence. The trimerization domain (TrD) is shown in red, the pleckstrin homology domain (PHD) in orange, and the fusion domain (FD) in yellow. Domains are connected by segments (named R1 to R5) that refold during conformational change: segments R1 and R4 are in cyan, segments R2 and R3 are in green, R5 and the CTD are in light and deep purple, respectively. Transmembrane domain (TMD) and intra-cytoplasmic domain (ID), which are not resolved in our structures, are also indicated. **b-c** Cryo-EM density map of VSV G incubated at pH 8.0 (**b**) and at pH 5.5 (**c**). G ectodomain is in skyblue, the LMNG micelle is in grey, and starting residues of the glycosylation chains are in cyan. The part of the fusion domain inserted in the micelle is colored in yellow, and the CTD in magenta. **d-e** Close-up view of the fusion loops insertion into the micelle at pH 8.0 (**d**) and pH 5.5 (**e**). The outline of the micelle is indicated by the red dotted line. The hydrophilic residues located in the hydrophilic shell are depicted in green. The hydrophobic residues in the fusion loops are in orange.

To date, no complete structure (*i.e.* including the TMD and the intra-viral or cytoplasmic domain -CD-) of VSV G has been reported. It should also be noted that only part of the structure of the CTD is known (up to residue 432 for the pre-fusion state and residue 410 for the post-fusion state). Interestingly, several mutagenesis studies suggest that the CTD, along with the TMD, are critical for G to mediate membrane fusion, especially in the final steps of the process ^25–28^. Thus, the mechanistic steps of G refolding and membrane fusion process are not fully understood due to the lack of structural information on the CTD and TMD.

VSV G ectodomain is the primary target of neutralizing antibodies. Although the antigenicity of VSV G has been characterized ^29,30^, no structural data of any VSV G / monoclonal antibody (mAb) complex have been reported so far.

Here, we report a cryo-electron microscopy (cryo-EM) study of detergent-solubilized full-length VSV G purified from viral particle. This study allows us to complete the structures of the ectodomain in its pre- and post-fusion conformations in a more physiological context. In particular, we extend our knowledge of the structure of the CTD in the post-fusion conformation and provide data that indicate that the TMDs are not visible in the detergent micelle, suggesting that they are rather mobile within the membrane. We also describe the first structures of VSV G bound to a commercial neutralizing antibody (mAb 8G5F11). This mAb binds G in both its pre- and post-fusion conformations. These first structures of a complex between G and a neutralizing antibody allowed the characterization of the epitope in detail and the identification of key residues on VSV G involved in this interaction. Globally, this work increases our knowledge of the structure of VSV G, the most widely used viral glycoprotein for the delivery of cargo and in gene therapy for lentivirus pseudotyping, and its low-pH induced refolding pathway.

## Results

### Overall structures of detergent solubilized VSV G in the pre- and post-fusion states

To gain insight into the full-length structure of VSV G *i.e* the ectodomain plus the TMD and CD (**Fig. 1a**), we developed a protocol to isolate VSV G (Indiana strain) from concentrated virus particle preparations without any affinity purification tag. Briefly, VSV G was extracted directly from viral membranes using lauryl maltose neopentyl glycol (LMNG) and subsequently purified to homogeneity through successive chromatography steps (**Supplementary Fig. 1**). We then performed cryo-EM single-particle analysis (SPA) on VSV G incubated at pH 8.0 (VSV G pH 8.0) and at pH 5.5 (VSV G pH 5.5) (**Supplementary Fig. 2-3, Table S1**).

2D class averages were generated for both VSV G pH 8.0 and VSV G pH 5.5. Analysis of the 2D classes revealed a preferential orientation of the molecule in both conditions, with approximately 80 % of the particle exhibiting top views (**Supplementary Fig. 2-3**). Using automated particle picking to enrich the data set with sides views, data processing yielded reconstructions with an overall resolution of 2.9 Å for both the high- and low-pH structures (**Fig. 1b-c**).

The overall structure of the ectodomain in both reconstructions was consistent with previous studies, showing a trimeric spike with dimensions in height of 8 nm at pH 8.0 and 12 nm at pH 5.5, corresponding to the pre- and post-fusion conformations ^8,31^, respectively (**Fig. 1b-c**) and associated with a detergent micelle at the FD tips. Notably, the presence in the sample of TMD and CD did not appear to influence the global structure of VSV G ectodomain (**Fig. 1b-c**). Indeed, structural overlay of the crystallographic- and cryo-EM-derived models yielded root-mean-square deviations (RMSD) for the Cα backbone of 0.78 Å for the pre-fusion structure and 0.69 Å for the post-fusion structure (PDB code 6TIT^15^ and 5I2M ^16^ respectively) (**Table S2**). Additionally, there was a very clear density for the starting residues of the two glycosylation chains (attached to N163 and N325) (Machamer and Rose, 1988) in both structures (**Fig. 1b-c**). It is worth noting that comparison with the crystallographic structure of the post-fusion state of VSV G ^16^ revealed an additional density corresponding to part of the CTD in the ectodomain of the post-fusion cryoEM structure (**Fig. 1c**, in magenta).

Despite repeated attempts to visualize the content of the detergent micelle using several computational cryo-EM methods (such as 3D-classification, 3DVA, 3DFLEX and several masking strategies), the TMDs (resp. the CDs) were not discernible within (resp. in the vicinity of) the micelle (**Fig. 1b-c**) in both conformations, although SDS-PAGE did not reveal any proteolysis (**Supplementary Fig. 1**).

On the other hand, the tips of FDs were visible inside the micelles, suggesting that the absence of detection of TMDs in the density is due to the fact that TMDs are mobile and do not have a fixed position within the micelle. Measurements indicate that the penetration depth of FDs into the micelle is approximately 9 Å for both conformations (**Fig. 1d-e**). This is consistent with the organization of the micelle of LMNG having a hydrophobic core radius of 14 Å and a hydrophilic shell thickness of 7 Å ^32^. This allows the accommodation of the hydrophobic residues in the fusion loop in the hydrophobic core as well as several polar or charged residues in the hydrophilic shell (Fig. 1d-e).

### Reorganization of the C-terminal linker region

The resolution of the CTD in the post-fusion conformation is approximately 4.5 Å, allowing for its partial tracing up to residue 426 (**Fig. 2a**). For each protomer, the traced segment extends in a predominantly hydrophilic groove (**Fig. 2b**) located between this protomer (protomer A) and the neighboring one (protomer B). Structural analysis reveals a hydrophobic patch on the CTD, composed of residues L423 and F425, which interact closely with residue I82 located on the FD of the same protomer (protomer A) and residue L105 and P107 of the FD of the neighboring protomer (protomer B) (**Fig. 2c**). This hydrophobic interaction stabilizes the C-terminal domain against the FD. Sequence alignment of Vesiculovirus glycoproteins indicates that the hydrophobic nature of these residues is conserved among the genus (**Supplementary Fig. 4**).

**Fig. 2.**
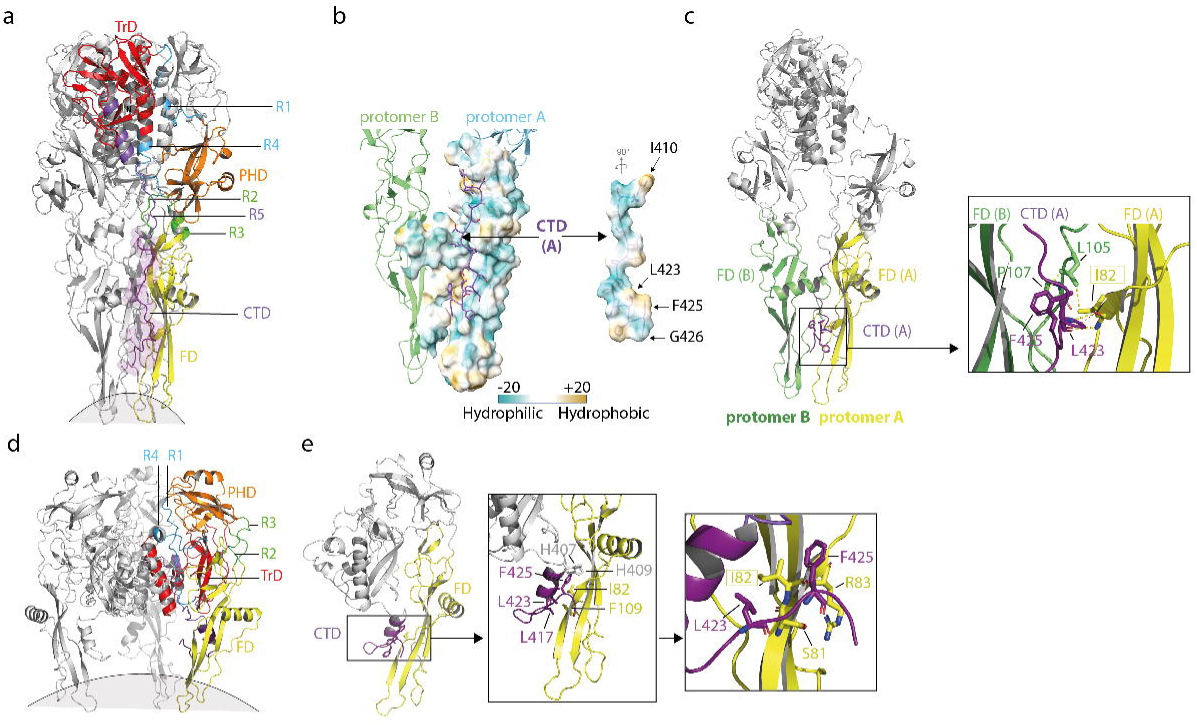
VSV G CTD rearrangement. **a** Ribbon representation of VSV G trimer in its post-fusion conformation, fitted and traced up to residue 426 in the cryo-EM map obtained at pH 5.5. The same color code as in Fig. 1a is used. **b** Hydrophobic map of the surfaces of interaction between segment 410-426 of chain A (right) and neighboring FD (left) in the post-fusion conformation. The hydrophobicity scale is indicated. **c** The hydrophobic patch at the level of the CTD (L423 and F425) interacting with I82 and L105 stabilizes the CTD close to the FD in the post-fusion conformation. **d** Ribbon diagram of VSV G trimer in its pre-fusion conformation, fitted and constructed in the cryo-EM map obtained at pH 8.0. The same color code as in Fig. 1a is used. **e** Close-up views on VSV G CTD in the pre-fusion conformation. The middle panel shows the folding of the CTD. The left panel provided a detailed view of the environment surrounding L423 and F425 in VSV G pre-fusion conformation (left panel).

In the pre-fusion conformation, the overall structure of the ectodomain was similar to previously determined crystal structures ^15^, with the CTD modeled up to residue 426 (**Fig. 2d-e**). In both X-ray crystallography and cryo-EM pre-fusion structures, the CTD forms a small helix (res 409 to 416) followed by a β-turn between P418 and E421 and a short β-hairpin (res 422 to 432, only visible up to res 426 in the cryo-EM structure) forming a hook-like structure (**Fig. 2e**). This indicates that the presence (or absence) of the TMD of VSV G does not interfere with its fold. Notably, in the pre-fusion conformation, the hydrophobic residues L423 and F425 on the CTD also form a hydrophobic cluster with I82 on the FD (**Fig. 2e**). Thus, this hydrophobic cluster is present in both the pre- and post-fusion states and, in both cases, it locks the CTD in close proximity to the FD (**Fig. 2c and Fig. 2d-e**). However, the organization of the CTD is completely different between the two structures (**Fig. 2c and Fig. 2d-e**). Thus, the CTD, as R1, R2, R3, R4 and R5 segments, refolds during the low-pH induced structural transition of VSV G. In fact, the CTD and R5 are part of a single long segment that refolds during G conformational change.

### Neutralization of VSV by mAb 8G5F11

The commercial monoclonal antibody 8G5F11 ^29^ neutralizes VSV and other Vesiculoviruses ^30^. To characterize the interaction of VSVG with mAb 8G5F11, we incubated purified mAb at various pH values with magnetic beads coated with protein A, followed by the addition of purified full-length VSV G pre-incubated at pH values between 5.0 and 8.0. After an incubation of 30 min, beads were washed, and the associated proteins were analyzed by SDS-PAGE. These interaction experiments demonstrated that 8G5F11 is able to bind VSV G over a range of pH values from 8.0 to 5.0 (**Fig. 3a**). This suggested that 8G5F11 recognizes several VSVG conformations.

**Fig. 3.**
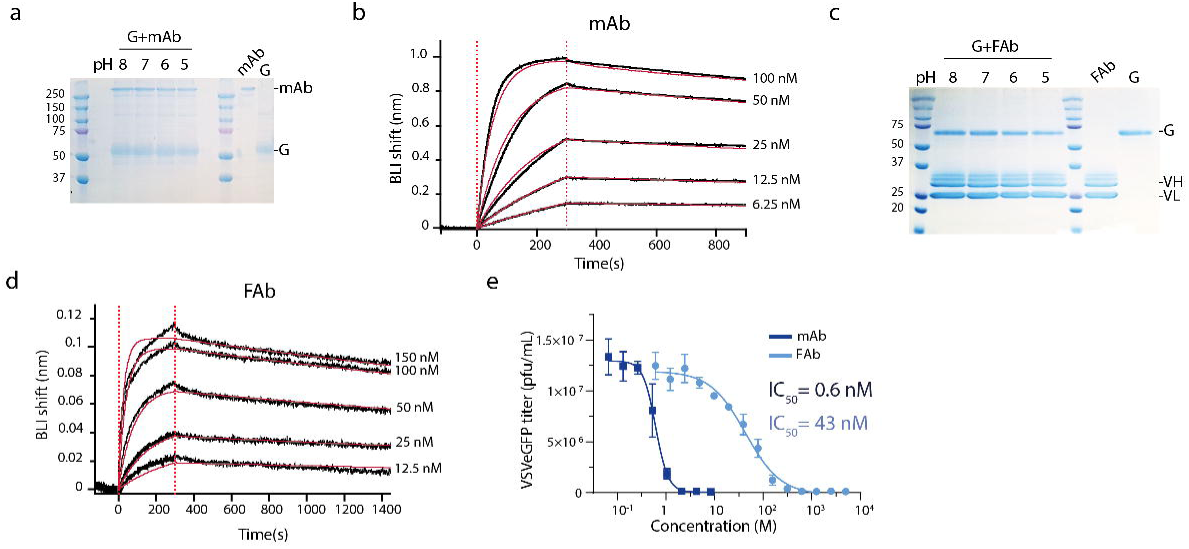
mAb 8G5F11 interacts with the pre- and the post-fusion conformations of VSV G. **a** Coomassie-stained SDS-PAGE analysis (non-reducing conditions) of interaction experiments between VSV G and mAb 8G5F11 bound to protein A coated magnetic bead, incubated at various pHs. **b** BLI sensograms showing the binding kinetics of VSV G_ect_ to mAb 8G5F11 at pH 8.0. Experimental curves (black) were fitted (red) using a 1:1 binding model. One representative experiment over 3 is shown. **c** Coomassie-stained SDS–PAGE analysis (reducing conditions) of interaction experiment between VSV G and FAb 8G5F11 bound to strep-tactin coated magnetic beads, incubated at various pHs. **d** BLI sensograms depicting the binding kinetics of VSV G_ect_ to 8G5F11 FAb at pH 8.0. Experimental curves were fitted using a 1:1 binding model. One representative experiment over 3 is shown. **e** Neutralization of VSVe-GFP by mAb (dark blue) and FAb (light blue) by 8G5F11 mAb (left panel) and FAb (right panel). VSV-eGFP was preincubated with increasing concentrations of mAb or Fab. At 5 hours *p.i.,* the percentage of infected cells determined by counting the number of cells expressing eGFP using flow cytometry. This was used to calculate the infectious viral titer. Data depict the mean with standard error from experiments performed in triplicate. Average IC_50_ value are indicated.

Additionally, we employed bio-layer interferometry (BLI) to investigate the binding parameters of coated mAb 8G5F11 to VSV G_ect_ at pH 8.0. For this, we used the thermolysin-generated VSV G_ect_ (aa residues 1–422) that is monomeric in solution ^8^. Analysis of the BLI data revealed that mAb 8G5F11 exhibited a dissociation constant in the nanomolar range for VSV G_ect_ at pH 8.0 (**Fig. 3b**).

We engineered the corresponding FAb fragment of mAb 8G5F11. For that, variable heavy (VH) and light (VL) chains of mAb 8G5F11 were sequenced and cloned into an insect cell expression vector (**Supplementary Fig. 5**). The construction resulted in a FAb C-terminally fused to a strep-TagII. After establishment of the stable S2 cell line, recombinant FAb was purified from cell culture media by affinity chromatography followed by SEC (**Supplementary Fig. 5**). Interaction experiments conducted under the same conditions as for the mAb 8G5F11 (except that the magnetic beads were coated with strep-tactin) show that FAb 8G5F11 binds VSV G across the same pH range, with BLI measurements also indicating a binding affinity in the nanomolar range (**Fig. 3c-d**).

To determine the IC_50_ values of mAb 8G5F11 and its corresponding FAb, we performed neutralization experiments by pre-incubating a recombinant VSV that express eGFP (VSV-eGFP) with serial dilutions of mAb or FAb. After 30min of incubation, the inoculum was transferred onto HEK-293T cells. After 5h of infection, the percentage of infected cells (expressing eGFP) was measured by flow cytometry. Both mAb 8G5F11 and FAb 8G5F11 inhibit VSV infection in a dose-dependent manner. Calculated viral titers were plotted against the concentration of mAb or FAb and the dose-response curve was fitted using the Hill equation (**Fig. 3e**). The estimated IC_50_ value for the FAb 8G5F11 was approximately 40 nM, whereas it decreased to 0.6 nM when neutralization was performed with the mAb 8G5F11. FAb 8G5F11 neutralization curve exhibited a Hill coefficient of 1.2±0.17 whereas mAb neutralization curve had a Hill coefficient of 3.35±0.79 indicating some cooperativity effect (**Fig. 3e**). This is consistent with the divalent nature of the mAb, which induces avidity-enhanced binding to VSV G, as well as with its larger size, which increases steric hindrance, potentially decreasing the number of mAbs required to neutralize a virion compared to that of FAbs.

### Structural characterization of FAb 8G5F11 bond to VSV G

To better understand how mAb 8G5F11 binds VSV G at different pH values, we performed cryo-EM SPA on VSV G in complex with FAb 8G5F11 at pH 8.0 and pH 5.5 (**Supplementary Fig. 6-7, Table S1**). The cryo-EM structure of VSV G in complex with the FAb revealed a stochiometric binding of one FAb per protomer at both pHs (**Fig. 4a** and **Fig. 4c**). At pH 8.0, the FAb binds each VSV G protomer at the top of the molecule (**Fig. 4a-b**) with an angle of approximately 45° with respect to the axis 3 of the trimer (**Fig. 4b**). At pH 5.5 the binding site is the same but the relative orientation of the FAb is different with an angle of 135° (**Fig. 4d**).

**Fig. 4.**
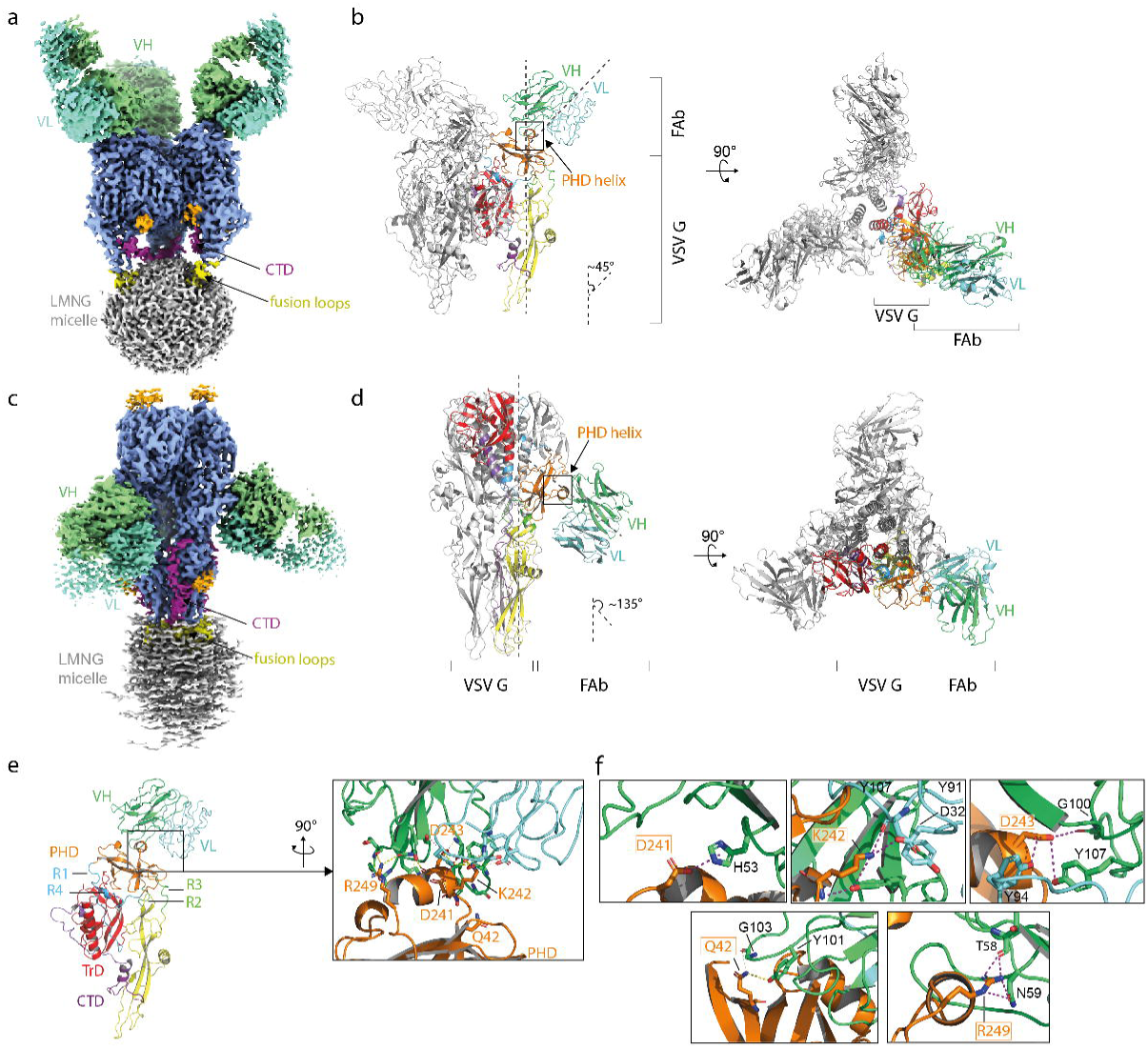
Structural basis for the neutralization of VSV by mAb 8G5F11. **a** Cryo-EM density map of VSV G in complex with FAb 8G5F11 at pH 8.0 shown in side view orientation. One 8G5F11 FAb molecule was observed to bind to each protomer of VSV G. G ectodomain is in sky-blue, the LMNG micelle in grey, the starting residues of the glycosylation chains are in orange. Fusion loops are in yellow and CTD in magenta. FAb chains are in green and cyan. **b** Ribbon diagram of VSV G trimer in the pre-fusion conformation in complex with FAb 8G5F11, viewed from the side (left panel) and from the top (right panel). The VSV G model was fitted in the density and traced up to residue 426. The variable heavy (VH) and light (VL) chains of the 8G5F11 Fab were modelized and fitted in the identified density. The same color code for G as in Fig. 1a is used. **c** Cryo-EM density map of VSV G in complex with FAb 8G5F11 at pH 5.5, shown in side orientation. The same color code as in Fig. 4a is used. **d** Ribbon diagram of VSV G post-fusion trimer in complex with FAb 8G5F11, viewed from the side (left panel) and from the top (right panel). As in the pre-fusion state, one molecule of FAb binds each VSV G protomer of the post fusion trimer. The VSV G model was fitted in the density and traced up to residue 426, while the variable heavy (VH) and light (VL) chains of the 8G5F11 Fab were fitted in the identified density. The same code for G as in Fig. 1a **is used**. FAb VH chain is in green and VL chain is in cyan. **e-f** Interaction between 8G5F11 and VSV G. Key residues are depicted in stick representation and are highlighted in the boxed panels in (**f)**. **f** Close up on the residues involved in the antibody-antigen interaction at high pH. VSV G PHD is in orange, and the key residues interacting with 8G5F11 are depicted in sticks.

The binding of the FAb did not influence the structure of the ectodomain; at pH 8.0 VSV G is in its pre-fusion conformation and at pH 5.5 VSV G is in its post-fusion conformation. The complexes between G and the FAb did not allow us to determine the structure of new parts of G.

In both structures, the interface between the FAb and VSV G is the same and principally involves the variable heavy chain (VH) of the FAb binding to the VSV G PHD **(Fig. 4b, 4d and 4e)**. The interaction is mainly hydrophylic. Three VSV G residues (D241, K242, and D243) are located at the core of the interface between G and 8G5F11 paratope. The carboxylic group of D241 establishes polar contact *via* its side chain with residue H53 of the FAb. The side chain of D243 establishes a cluster of polar contacts with the hydroxyl groups of Y94 (on the VL chain) and Y107 (on the VH chain) as well as with the carbonyl of the main chain of G100 (on the VH chain). VSV G K242 side chain interacts with the carboxyl group of D32 and the main chain of Y91 (on the VL chain) and the carbonyl group of K242 interacts with the hydroxyl group of Y107 (on the VH chain). Two additional VSV G residues (Q42 and R249) located outside of the core also contribute to the interaction. The amide group of side chain of residue Q42 establishes hydrogen bonds with the carbonyl of the main chain of G103 and the hydroxyl group of Y101 on the VH chain of the FAb. Finally, guanidine group of R249 on VSV G establishes H-bonding with N59 side chain and the T58 main chain (both on the VH chain).

### Characterization of 8G5F11 binding site

To investigate the contribution of residues constituting the mAb 8G5F11 binding site on VSV G (**Fig. 4e**), we generated alanine mutants at positions Q42, D241, K242, D243, R249 of G. HEK-293T cells were transfected with plasmids encoding either wild type (WT) VSV G or VSV G alanine mutants (G_mut_: Q42A, D241A, K242A, D243A or R249A). Surface expression of the VSV G mutants was confirmed using a conformational probe, GST-CR3 (glutathione S-transferase fusion with the CR3 domain of the LDL-R ^2^), labeled with a fluorescent dye ATTO^550^ (**Fig. 5a**). Indeed, GST-CR3 binds VSV G in its pre-fusion conformation and its binding site ^2,15^ is distinct from the binding site of mAb 8G5F11 (**Supplementary Fig. 8**). Flow cytometry analysis confirmed that all mutants were correctly expressed at the cell surface, as evidenced by their efficient binding to the CR3-GST conformational probe (**Fig. 5b**). Then, we assessed the ability of VSV G mutants to bind mAb 8G5F11 and its corresponding FAb on the cell surface. Mutations D241A and D243A induced a significant reduction in binding for both the mAb and FAb 8G5F11 (**Fig. 5c**). Afterwards, we evaluated whether these mutant glycoproteins (VSV G_D241A_ and VSV G_D243A_) could maintain viral infection. We employed a recombinant VSV (VSVΔG) lacking the G envelope gene, replaced by the green fluorescent protein (GFP) gene pseudotyped with VSV glycoprotein (**Fig. 5d**). The incorporation of VSV G mutants into the membrane of VSV pseudotypes was verified by Western-blot analysis (**Fig. 5e**). Indeed, the VSVΔG/G_D241A_ and VSVΔG/G_D243A_ pseudotypes exhibit an equivalent level of glycoprotein incorporation to that observed with VSVΔG/G_WT_. Infectivity of VSV pseudotypes in HEK-293T cells was determined by flow cytometry, measuring the percentage of infected cells 16 hours *p.i.* In the absence of antibody, both VSVΔG/G_WT_ and VSVΔG/G_mut_ pseudotypes exhibited comparable infectious titers, approximately 2 x 10^7^ pfu/ml (**Fig. 5f**, left panel). We then performed neutralization experiments by pre-incubating VSVΔG/G_WT_ and VSVΔG/G_mut_ inoculum with serial dilutions of mAb or FAb 8G5F11 (**Fig. 5f**). As expected, VSVΔG/G_WT_ pseudotype was efficiently neutralized by mAb and FAb 8G5F11, with IC_50_ values of 0.2 nM and 10 nM respectively (**Fig. 5f**, right panel). In contrast, VSVΔG/G_D241A_ and VSVΔG/G_D243A_ pseudotypes were not neutralized by FAb or mAb 8G5F11 at either concentration, indicating that these residues play an important contribution to mAb 8G5F11 binding and neutralization activity (**Fig. 5f**, left panel). It is noteworthy that the acidic character of the residues corresponding to D241 and D243, is widely conserved among vesiculoviruses (**Supplementary Fig. 4**). This explains why 8G5F11 is broadly neutralizing ^30^.

**Fig. 5.**
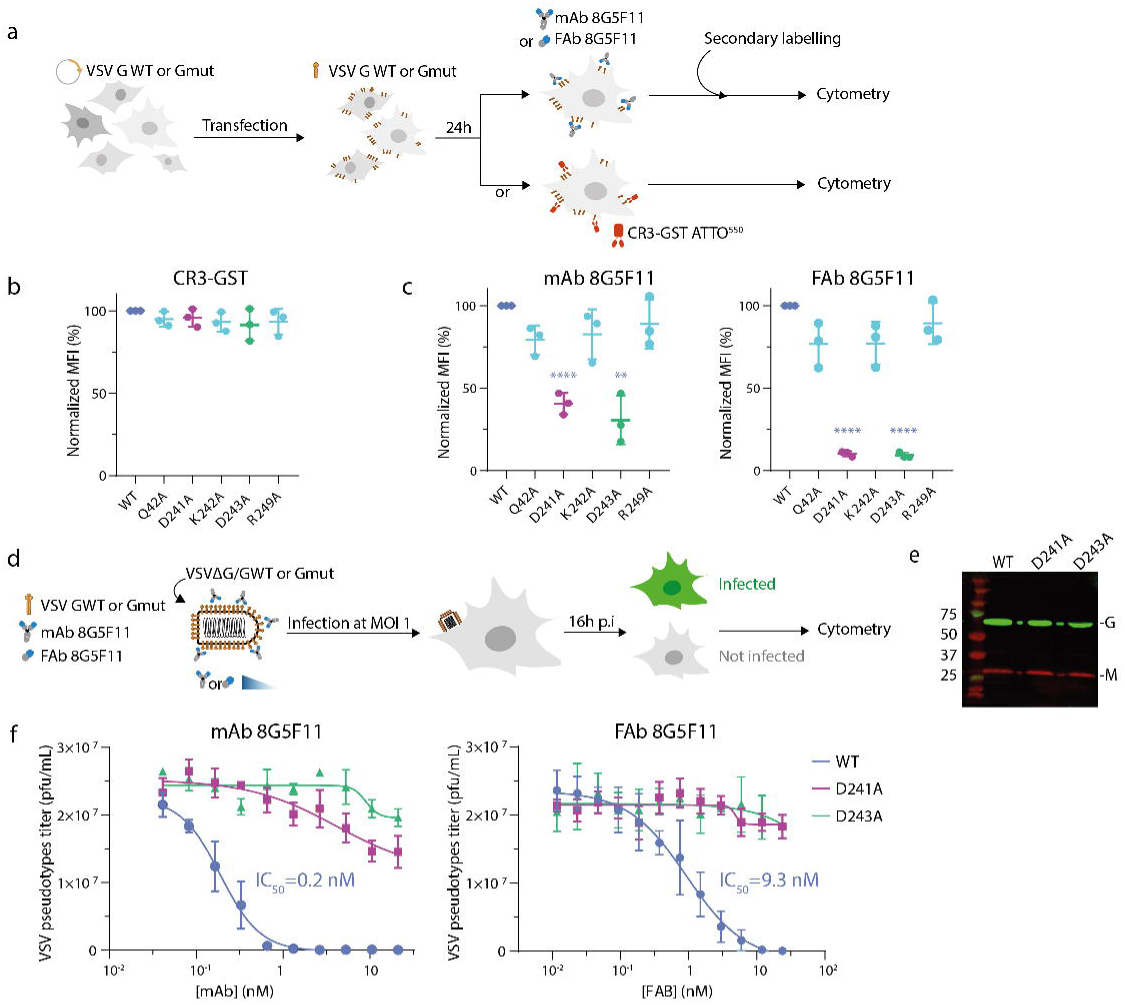
Residues D241 and D243 are essential for the interaction of 8G5F11 with VSV G and for neutralization. **a** Schematic description of the binding assays used to assess the capacity of VSV G WT and VSV G alanine mutants to bind CR domains and 8G5F11 mAb or FAb. **b** Surface expression test for VSV G WT and VSV G alanine mutants by measuring their ability to bind the conformational probe (CR3-GST labelled by and ATTO^550^ fluorophore). The histogram indicates the mean fluorescence intensity of ATTO^550^ positive cells for each G construct. Three independent experiments were performed. Error bars represent the standard deviation. **c** Antibody recognition assay for VSV G WT and mutants G by measuring their ability to bind 8G5F11 mAb (left panel) or FAb (right panel). **d** Schematic description of the neutralization assay using VSVΔG/G pseudotypes. **e** Incorporation of VSV G WT and VSV G alanine mutants in VSVΔG viral particles, assessed using a polyclonal anti-VSV G and an anti-VSV M antibodies. **f** Neutralization of VSVΔG/GWT, VSVΔG/GD241A, and VSVΔG/GD243A by mAb (left panel) or FAb 8G5F11 (right panel). VSV pseudotypes were preincubated with increasing concentrations of mAb or Fab. At 16 hours *p.i.,* the percentage of infected cells was determined by counting the number of cells expressing eGFP using flow cytometry. This was used to calculate the infectious viral titer. Data depict the mean with standard error for experiments performed in triplicate. Average IC_50_ are indicated.

## Discussion

In this work, we determined the structures of full-length VSV G, solubilized from viral particles, using cryo-EM SPA, both alone and in complex with the FAb derived from the neutralizing antibody 8G5F11. Overall, these structures of G are largely consistent with those obtained by crystallography ^14–16^ and correspond to the pre- and post-fusion conformation of VSV G. However, the cryo-EM structures of the post-fusion state of VSV G reveal new details about the organization of the CTD that were absent in previously reported post-fusion crystal structures of Vesiculovirus Gs ^16,21^. Since the CTD organization in the pre-fusion conformation was already known ^15^ and confirmed through the new cryo-EM structures of the pre-fusion form, this work shows that this domain also undergoes a major conformational change with a complete reorganization of secondary structures and a relocalization in a hydrophilic groove between two adjacent FDs. This organization of the CTD locks the post-fusion conformation and contributes to the stability of the post-fusion trimer. The correct positioning of the CTD within the hydrophilic groove is ensured by the hydrophobic patch formed by residues L423 and F425 with residue I82 of the same protomer and residues L105 and P107 of the adjacent protomer. The hydrophobic character of these residues is conserved among all Vesiculoviruses.

Previous studies have shown that the low pH-induced conformational change is initiated by refolding of the CTD ^15,22^, which dissociates from the rest of the protomer, allowing the exposure of fusion loops at the top of the molecule, which enables its interaction with the target membrane. This work shows that the repositioning of the CTD that stabilizes the post-fusion state occurs at a late stage of the refolding process in agreement with the crystal structures of intermediate conformations on the transition pathway of CHAV G and RABV G ^22,33^.

Despite significant efforts, we could not visualize the TMD and CD in the cryo-EM densities. This is in striking contrast to the fact that we were able to visualize the part of the FD inserted into the detergent micelle. This suggests that the TMD exhibits a high degree of mobility in the micelle and probably also in the membrane. This contrasts with the herpesvirus gBs for which the TMD is visible in the post-fusion structure of HSV1 gB ^34^ and in the pre-fusion structure of gB of HCMV ^35^. It should be noted that the C-terminal part of the CTD corresponding to residues 432-446 for VSV G (missing in cryo-EM structures) is rather short compared to the corresponding membrane proximal external region (MPER) of gBs which is approximately 50 residues long and forms a flat-lying helical structure on the micelle that may contribute to stabilization of TMD orientation.

It has been suggested that flexibility between the ectodomain and the TMD is required for the proper refolding of viral fusion glycoproteins ^36^. This flexibility was demonstrated for the hemagglutinin (HA) of the influenza virus ^36^. However, in this case, despite the flexibility of HA, the trimeric organization of TMDs was visualized after masking the ectodomain. In contrast, using a similar approach for VSV G, we were unable to resolve its TMDs. Thus, this raises the possibility that VSV G TMDs may not associate into a trimeric structure. Supporting this hypothesis, tomographic reconstructions of VSV particles, showed no density potentially corresponding to TMDs ^37^. Similarly, in the cryo-EM structure of rabies virus G (also solubilized with LMNG), the TMDs were indistinguishable, likely due to their high level of flexibility ^38^.

Additionally, our work also provides the structures of VSV G in complex with FAb 8G5F11. VSV Indiana neutralizing mAb 8G5F11 cross-react with the glycoprotein of other vesiculoviruses including Maraba, Alagoas, Cocal, and New Jersey ^30^. The structures revealed that the epitope recognized by the mAb corresponds to a helix of the PHD that is exposed at the top of G protomers in their pre-fusion conformation. This region corresponds also to the major antigenic site (major antigenic site II) of RABV ^39,40^. Three G residues (D241, K242, D243) from the core of the interface between G and the paratope, among which residues D241 and D243, are crucial for mAb binding and neutralization. Acidic residues in position 241 and 243 are widely conserved among Vesiculoviruses neutralized by 8G5F11, but they are absent in Piry virus G, which is not neutralized ^30^, as well as in Isfahan virus and CHAV Gs. It should be noted that CHAV G has not the same structural organization in this region: instead of a helix, it has a small β-hairpin ^21^.

The molecular mechanisms underlying this neutralization have not been investigated. However, it is plausible that the binding of the antibody creates a protein layer around the virion, increasing the distance between the virus and the cellular membrane, thus impeding receptor recognition (although the antibody does not bind directly to the receptor-binding site). Alternatively, it may interfere with the conformational changes required for membrane fusion (due to the proximity of the spikes in the viral membrane) or with the formation of the helical network of spikes in their post-fusion conformation that is supposed to play a role in the late stages of the fusion process ^31^.

Although VSV and related viruses do not pose a major threat for human health, outbreaks in horses and livestock can cause economic losses. A better understanding of VSV neutralization may lead to the conception of vaccines protecting against a broad spectrum of vesiculoviruses. Furthermore, VSV is a promising candidate for oncovirotherapy due to its ability to selectively replicate in and kill cancer cells ^41–43^. However, it still has the potential to spread uncontrollably in the bodies of patients whose immune systems have been weakened by other anticancer therapies. The administration of neutralizing human or humanized monoclonal antibodies could offer a potential solution in such cases.

## Methods

### Viruses and cells

BSR, a clone of BHK21 (Baby Hamster Kidney cells; ATC CCCL-10) and HEK-293T (human embryonic kidney cells expressing simian virus 40T antigen; ATCC CRL-3216) were cultured in DMEM supplemented with 10% FCS, 50 U/mL of penicillin and 50 μg/mL of streptomycin at 37 °C in a humidified incubator with 5% CO2. Drosophila Schneider 2 (S2) cell line (Invitrogen) were grown in ESF 921 medium (Clinisciences) supplemented with 50 U/mL of penicillin and 50 μg/mL of streptomycin at 28° C.

Wild-type VSV MS (Mudd-Summer strain, Indiana serotype), VSVeGFP (a gift from Denis Gerlier) were propagated on BSR cells.

VSVΔG-GFP was propagated on HEK-293T cells that had been previously transfected with pCAGGS VSV G.

### Plasmid and cloning

Point mutations were introduced starting from the cloned VSV G gene (Indiana Mudd-Summer strain; MS) in the pCAGGS plasmid. Briefly, forward and reverse primers containing the desired mutation were combined separately with one of the primers flanking the G gene to generate two PCR products. These two G gene fragments overlapped in the region containing the mutation and were assembled into pCAGGS vector,wich had been linearized using EcoRI, using Gibson assembly reaction kit (New England Biolabs).

### Purification of VSV G protein

VSV G protein was directly solubilized from concentrated stocks of VSV MS using 2% w/v laurylmaltose neopentyl glycol (LMNG; Anatrace) for 22 hours at 37°C. Insoluble material was removed by centrifugation (20 minutes at 20 000g, 4°C). The supernatant was then loaded onto an anion exchange chromatography column (ResourceQ, Cytiva) equilibrated with 20mM Tris-HCL pH 8.0, 2mM EDTA, 0.1 % w/v LMNG. VSV G was eluted with a linear gradient of NaCl (from 0 to 0.5 M NaCl). An additional purification step was performed on a size exclusion chromatography column (Superdex S200 increase column HR10/300, Cytiva) equilibrated in 20 mM Tris-HCl pH 8.0, 150 mM NaCl, 2 mM EDTA, 0.005 % w/v LMNG. Purified VSV G was concentrated to 5 mg/ml (Amicon Ultra 30kDa cut of; Millipore) and stored at -80°C.

### Production of recombinant FAb

The heavy and light chains (HC and LC respectively) of 8G5F11 mAb were sequenced (Absoute antibody; https://absoluteantibody.com/) and then cloned into an insect cell expression vector derived from the pMT-puro plasmid (Addgene #17923) using Gibson assembly reaction kit (New England Biolabs), as described in ^44,45^. Briefly, the LC (light chain) is flanked by a metallothionein (MT) promoter and the Drosophila Bip secretion signal (BIP ss) at the N-terminus, with a poly A tail at the C-terminus. It is followed by the Fd portion of the heavy chain which is also flanked by MT promoter and BIP ss at the N-terminus, and a strep-tag II at the C-terminus, followed by a poly A tail (**Supplementary Fig. S5**).

### Preparation of cryo-EM grids and data collection

For VSV G samples, grids were prepared by applying 4µL of VSV G concentrated at 1.95mg/ml to negatively glow discharged (25mA, 45s) UltraFoil® Au grids 300 R1.2/1.3 (QUANTIFOIL®) prior to plunge freezing using a Vitrobot Mark IV (FEI) (20°C, 100% humidity, 5.5s blot time and 0 blot force). To obtain VSV G in the post-fusion conformation, MES 1 M pH 5.5 was added to reach a final concentration of MES of 100 mM, sample was added 30 min at 37°C just before freezing. The grids were pre-screened on a 200 kV Glacios and automated data collection was performed at the beamline CM01 of ESRF (Grenoble, France) on a 300kV Titan Krios equipped with a K3 direct electron detector, coupled to an energy filter (Bioquantum LS/967, Gatan Inc, USA) ^46^. 8,840 and 6,864 movies were recorded with EPU (Thermofisher) for the pre- and post-fusion of VSV G respectively, with a pixel size of 1.05 Å/pixels, a total dose of 40.4 e-/Å and a defocus range of -0.8 to -2.2 µm.

Cryo-EM grids of purified VSV G in complex with 8G5F11 FAb at pH 5.5 were prepared by applying 3.5 µL of the complex at 0,83 mg/mLto negatively glow discharged (25mA, 45s) UltraFoil Au grids 300 R1.2/1.3 (QUANTIFOIL®) prior to plunge freezing using a Vitrobot Mark IV (FEI) (30°C, 100% humidity, 5.5 s blot time and blot force 0). The grids were pre-screened on a 200 kV Glacios electron microscope at IBS (Grenoble, France) and automated data collection was performed at the beamline CM01 of ESRF (Grenoble, France) on a 300 kV Titan Krios equipped with a K3 direct electron detector, coupled to an energy filter (Bioquantum LS/967, Gatan Inc, USA) ^46^. 19,423 movies were recorded with EPU (Thermofisher) for the pre- and post-fusion VSV G full length respectively, with pixel size of 0,84 Å/pixels, a total dose of 49,71 e-/Å and a defocus range of -0.8 to -2.2 µm.

Cryo-EM grids of VSV G in complex with 8G5F11 FAb at pH 8,0 were prepared by applying 4µL of the complex at a concentration of 0,79 mg/mL to negatively glow discharged (25mA, 45s) UltraFoil Au grids 300 R1.2/1.3 (QUANTIFOIL®) prior to plunge freezing using a Vitrobot Mark IV (FEI) (30°C, 100% humidity, 5.5 s blot time and blot force 0). SerialEM based automated data collection was performed a 200 kV Glacios electron microscope equipped with a K2 summit direct electron detector. 15,947 movies were recorded with EPU (ThermoFisher) with pixel size of 1.145 Å/pixels, a total dose of 38,5 e-/Å and a defocus range of -0.8 to -2.2 µm.

### Cryo-EM data processing and model building

Image processing was performed using cryoSPARC software ^47^ and is summarized in **Supplementary figures S2-3 and S6-7**. Recorded movies were dose-weighted and motion-corrected using cryoSPARC Patch Motion Correction program. CTF parameters were determined using cryoSPARC Patch CTF program. Particles were at first automatically picked using circular and elliptical blobs with a diameter between 100 and 230 Å for VSV G pH 5.5, between 100 Å and 250 Å for VSV G pH 8.0, between 150 Å and 300 Å for VSV G/Fab pH 8.0 and 80 Å and 300 Å for VSVG/FAb pH 5.0. A first round of particles picking was performed for each data set to select correct side views 2D classes. Topaz neural network was then used to enrich particles data set in particles side views and visible micelle ^48^. Several rounds of 2D classification were then performed in order to clean particle data sets. *Ab-initio* reconstruction was used to generate at least 2 models to classify particles among them. Several rounds of Homogeneous refinement and Non-uniform refinement ^49^ were performed without imposed symmetry. The best models were selected for global and local (per-particle) CTF refinement to improve particle, map quality and resolution. C3 symmetry was only imposed on VSV pH8.0 and VSV/Fab pH5.0 reconstructions. Reported maps resolutions were estimated by gold-standard Fourier shell correlation (FSC) of 0.143 criterion.

Model building for all protein chains was performed using COOT v.0.9.6 EL ^50^. Model refinement and validation was performed using Phenix v.1.20.1 ^51,52^ and ISOLDE ^53^. FAb model building was based on an initial protein model generated from AlphaFold2 ^54^ then fitted into the cryo-EM map and manually refined in COOT v.0.9.6 EL ^50^. Geometry was validated using MolProbity ^55^.

### Biolayer interferometry analysis (BLI)

BLI experiments were performed using an Octet® RED96e (Sartorius, Fremont, CA, USA) instrument operated at 20°C under 1000 rpm stirring with Octet BLI Discovery software (v.12.2.2.20). The 8G5F11 mAb was captured on Protein A-functionalized sensors at 20 µg/mL during 40 seconds in acetate 10 mM pH 4.0. FAb 8G5F11 were covalently immobilized with random orientation on Amine Reactive 2nd Generation sensors (AR2G) at 20 µg/mL during 600 seconds in acetate 10 mM pH 4.0 using standard EDC-NHs chemistry. Interactions with VSV G_ecto_, from 6.25 to 150 nM in running buffer (Tris-HCl 20 mM pH 8.0, NaCl 100 mM, EDTA 2 mM), were monitored during 300s (association phase). Dissociation step was set at 600s on the mAb surfaces and 1200 seconds on the FAb surfaces respectively. A bare sensor was always added as a reference surface to control the non-specific binding.

Data analysis was performed on Octet Analysis Studio software (v.12.2.2.26) and kinetics constant (ka and kdis) were extracted by fitting the data with a 1:1 binding model or a 2:1 heterogenous model, and equilibrium dissociation constant K_D_ was calculated.

The experiments were performed three times using similar values.

### Binding assay of 8G5F11 mAb and FAb to HEK-293T expressing GWT or Gmut at their surface

CR3-GST was produced in *E. Coli* (C41(DE3); Lucigen) and labeled with the fluorescent dye ATTO^550^ NHS ester (Sigma Aldrich) following the instruction of the manufacturer, as previously described (Nikolic et al., 2018). The labelled probe was then diluted at a concentration of 50 mM and stored at -80°C until use. The labelling ratio was estimated to be around 2 dyes per CR3-GST molecules.

Cell surface expression levels of G WT or mutant G were estimated by adding GST-CR3 ATTO^550^ at a concentration of 500 µM on transfected cells 24 hours post-transfection. The fluorescence of cells was measured using Beckman CytoFlex S cytometer.

For binding assays, HEK-293T cells were transfected with pCAGGS plasmids encoding G WT or G mutants using the calcium phosphate transfection method. Twenty-four hours after transfection, cells were collected and incubated with either 8G5F11 mAb (at a concentration of 1.3 10^-3^ mM) or 8G5F11 FAb (at a concentration of 2.5 10^-5^ mM) for 45 minutes at RT. Goat anti-mouse Alexa fluor 488 (at 1,3 10^-3^ mM) was added to the cells to accessed the binding of mAb 8G5F11. Anti-strep-tag coupled to PE fluorophore was added to the cells to accessed the binding of FAb 8G5F11. The fluorescence of cells was measured using a Beckman CytoFlex S cytometer.

### Virus neutralization assay

HEK-293T cells at 70% confluence in 96-well plates were infected at MOI 1 with VSV G-eGFP pre-incubated for 30 minutes with increasing concentrations of 8G5F11 mAb or FAb. 10^5^ VSVeGFP pfu were incubated with increasing concentrations of 8G5F11 Mab or Fab for 30 minutes. The percentage of infected cells (eGFP-positive) was determined using a Beckman CytoFlex S cytometer, 5 hours *p.i.*.

### VSVΔG-GFP pseudotypes neutralization assay

HEK-293T cells at 70% confluence were transfected with pCAGGS plasmids encoding G WT or mutant G using the calcium phosphate transfection method. Twenty-four hours after transfection, cells were infected with VSVΔG-GFP pseudotyped by VSV G WT at MOI 1. Two hours *p.i.*, cells were washed to remove the inoculum. Then, at 16 hours *p.i.*, supernatants containing the pseudotypes were collected. The infectious titers of the pseudotyped viruses was determined on non-transfected cells by counting cells expressing the GFP using a Beckman CytoFlex S cytometer 16 hours *p.i..* Experimental MOIs were precisely determined using the equation MOI=−ln[p(0)], where p(0) is the proportion of non-infected cells, and compared to that of VSVΔG-GFP pseudotyped with WT G which was normalized to 1.

### Western blot analysis

G WT and mutant G incorporation into the membrane of the pseudotyped particles was assessed, after concentrating viral supernatants, by Western blot analysis using P5D4, a mouse-monoclonal anti-VSV G antibody directed against the ICD of VSV G (1/5000) and a mouse-monoclonal anti-VSV M antibody (1/5000). Goat anti-rabbit DyLight 800 conjugate and Goat anti-mouse DyLigth 680 conjugate were used as secondary antibodies. Blots were imaged using a LI-COR Biosciences Odyssey®.

### Statistical analysis

All numerical data were calculated and plotted with mean ±SD resulting from three independent biological replicates. Results were analyzed using Prism v8.0.1 (GraphPad Software).

For the recognition assay, results are presented as normalized MFI. To do so, the geometric mean of the measured MFI for each condition of each experimental replicate was calculated. The geometric means were then normalized to the one of the WT. Results were compared using T-tests to compare means of biological replicates. Experiments were performed in triplicates.

For neutralization and pseudotypes assays, experimental titers were determined based on experimental MOIs. Dose-response parameters, namely Hillslope and IC_50_, were determined using a log(inhibitor) vs response – variable slope model.

## Acknowledgments

This work was supported by grants from the from the Agence Nationale de la Recherche, France, (ANR CE11GrLy) to AA. We acknowledge the cryoEM platform of I2BC, supported by the French Infrastructure for Integrated Structural Biology (FRISBI) (ANR-10-INSB-05-05). The present work has benefited from: the facilities and expertise of the I2BC platform PIM supported by French Infrastructure for Integrated Structural Biology (FRISBI) ANR-10-INBS-05. We acknowledge the European Synchrotron Radiation Facility for provision of beamtime on CM01 (proposal number MX-2440), and we thank A. Grinzato for assistance in using the beamline. The IBS/ISBG electron microscope facility is supported by the Auvergne-Rhône-Alpes Region, the Fondation Recherche Médicale (FRM), the fonds FEDER, and the GIS-473 Infrastructures en Biologie Santé et Agronomie (IBISA). This research used the platforms of the Grenoble Instruct center (ISBG; UMS 3518 CNRS-CEA-UGA-EMBL) with support from FRISBI (ANR-10-INSB-05-02) and GRAL within the Grenoble Partnership for Structural Biology (PSB).

## Author Contributions

A.A.A. and Y.G conceived the project. L.B. cultured and purified VSV. A.A.A., L.B. and A.A.A. initiated the project and performed early work, A.A.A., Y.G. and M.M. designed experiments. M.M. and LB performed experiments. M.O., L.Z. and G.S. collected the data. A.A.A., M.M performed reconstructions, built atomic models, and performed neutralization experiments, M.M., Y.G. and S.R. analyzed data. A.A.A. and Y.G. wrote the paper. M.M. S.R. and G.S. revised the manuscript. All authors read and approved the final version of the manuscript.

