## Supplementary figures legend for "Structures of vesicular stomatitis virus glycoprotein G alone and in complex with a neutralizing antibody"

**Supplementary information for “Structures of vesicular stomatitis virus glycoprotein G alone and in complex with a neutralizing antibody”**

Marie Minoves^1^, Malika Ouldali^1^, Laura Belot^1^, Stéphane Roche^1^, Lefteris Zakardas^2^, Guy Schoehn^3^, Yves Gaudin^1^, Aurélie Albertini^1^*

**Fig. S1 Purification of VSV G**

Elution profile of VSV G on a Superdex S200 HR 10/300 (cytiva) and Coomassie-stained SDS-PAGE analysis of VSV G purification steps (1: concentrated VSV preparation, 2: VSV G after anion exchange chromatography step, 3: VSV G after size exclusion chromatography step).

**Fig. S2 Cryo-EM data processing pipeline for VSV G at pH 8.0**

**a** Representative electron micrograph. **b** Cryo-EM data processing workflow in cryoSPARC. **c** Final density used for model building (left) and local resolution map calculated and plotted onto the sharpened VSV G reconstruction.

**Fig. S3 Cryo-EM data processing pipeline for VSV G at pH 5.5**

**a** Representative electron micrograph. **b** Cryo-EM data processing workflow in cryoSPARC. **c** Final density used for model building (left) and local resolution map calculated and plotted onto the sharpened VSV G reconstruction.

**Fig. S4 Sequence alignment of Vesiculovirus glycoproteins**

Conserved residues are highlighted in dark blue boxes, while similar residues are shown in lighter blue. VSVI G domains are indicated above the sequence and depicted according to Fig. 1a color code. Asparagine carrying N-glycosylation are marked with an orange circle. The residues constituting the fusion loops are framed in yellow boxes. Residues belonging to 8G5F11 epitope are framed in cyan boxes. The residues constituting the hydrophobic patch stabilizing the CTD in the post-fusion conformation are indicated by brown arrows.

**Fig. S5 Purification of VSV G-Fab complex for cryo-EM studies**

**a** Diagram of the construction used to produce 8G5F11 Fab. (BIP ss = BIP signal sequence; LC = light chain; Fd = part of the heavy chain composing the FAb; StrepII = strep-tag II).

**b** Elution profile of FAb on a Superdex S200 HR 10/300 (Cytiva) in 20 mM Tris-HCl pH 8.0, 150 mM NaCl 2 mM EDTA (left panel) and Coomassie-stained SDS-PAGE analysis of purified FAb.

**c** Schematic description of VSVG-FAb complex assembly for cryo-EM studies

**d** Elution profile of VSVG-Fab complex on a Superdex S200 HR 10/300 (Cytiva) in 20 mM Tris-HCl pH 8.0, 150 mM NaCl 2 mM EDTA (left panel) and Coomassie-stained SDS-PAGE analysis of purified VSVG-FAb.

**Fig. S6 Cryo-EM data processing pipeline for pre-fusion VSV G at pH 8.0 in complex with FAb**

**a** Representative cryo-electron micrograph.

**b** Cryo-EM data processing workflow in cryoSPARC.

**c** Final density used for model building (left panel) and local resolution map calculated and plotted onto the sharpened VSV G reconstruction (middle panel) and representative fit of atomic model of G and FAb into density (right panel).

**Fig. S7 Cryo-EM data processing pipeline for pre-fusion VSV G at pH 5.5 in complex with FAb**

**a** Representative cryo-electron micrograph.

**b** Cryo-EM data processing workflow in cryoSPARC.

**c** Final density used for model building (left panel) and local resolution map calculated and plotted onto the sharpened VSV G reconstruction (upper right panel) and representative fit of atomic model of G and FAb into density (lower right panel).

**Fig. S8 8G5F11 and LDL-R binding sites on VSV G**

Binding footprint of 8G5F11 and CR2 **(a)** or CR3 **(b)** on VSV G.
