## Supplementary figures and images for "Structures of vesicular stomatitis virus glycoprotein G alone and in complex with a neutralizing antibody"

### Supplementary Fig. 1

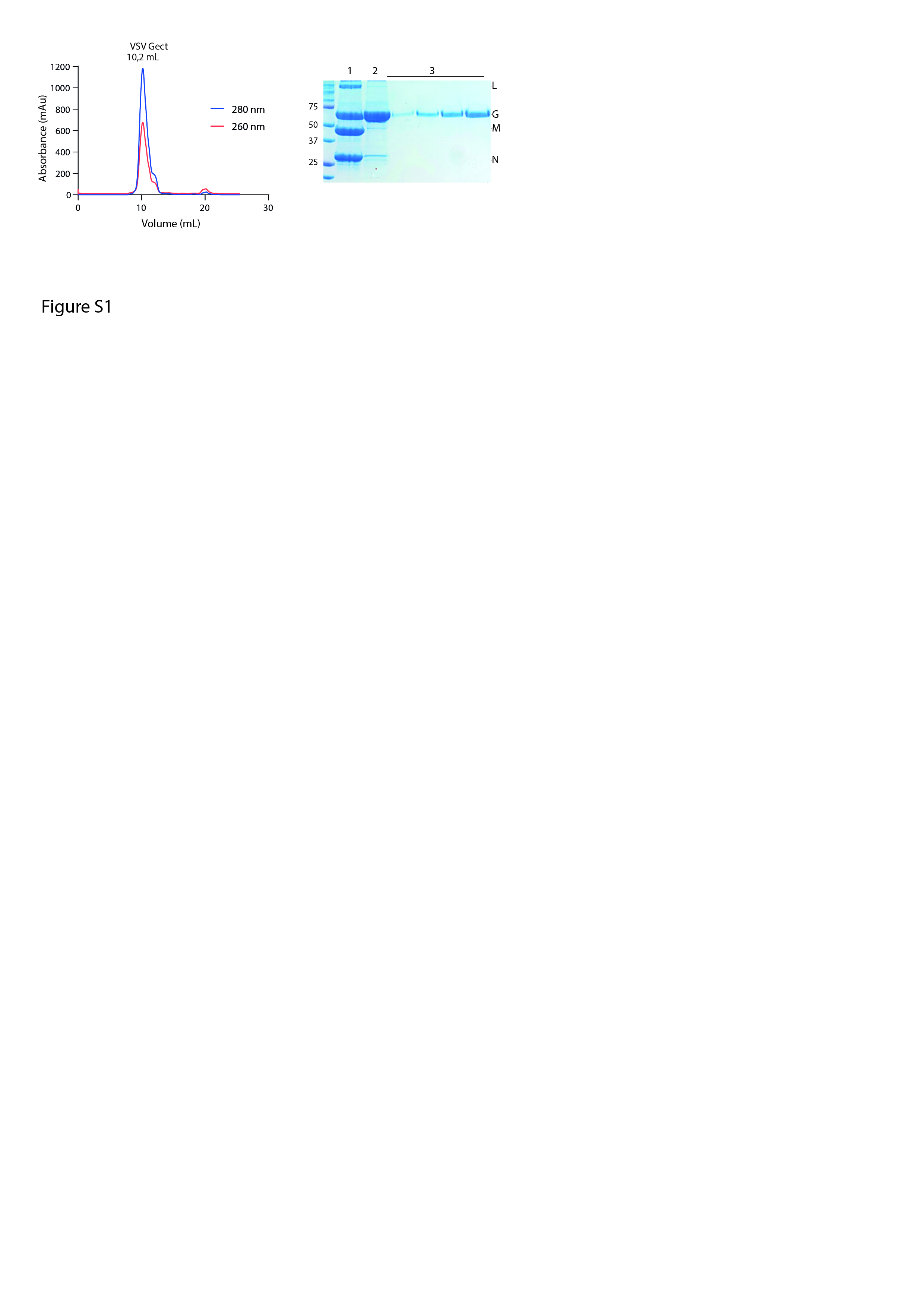

### Supplementary Fig. 2

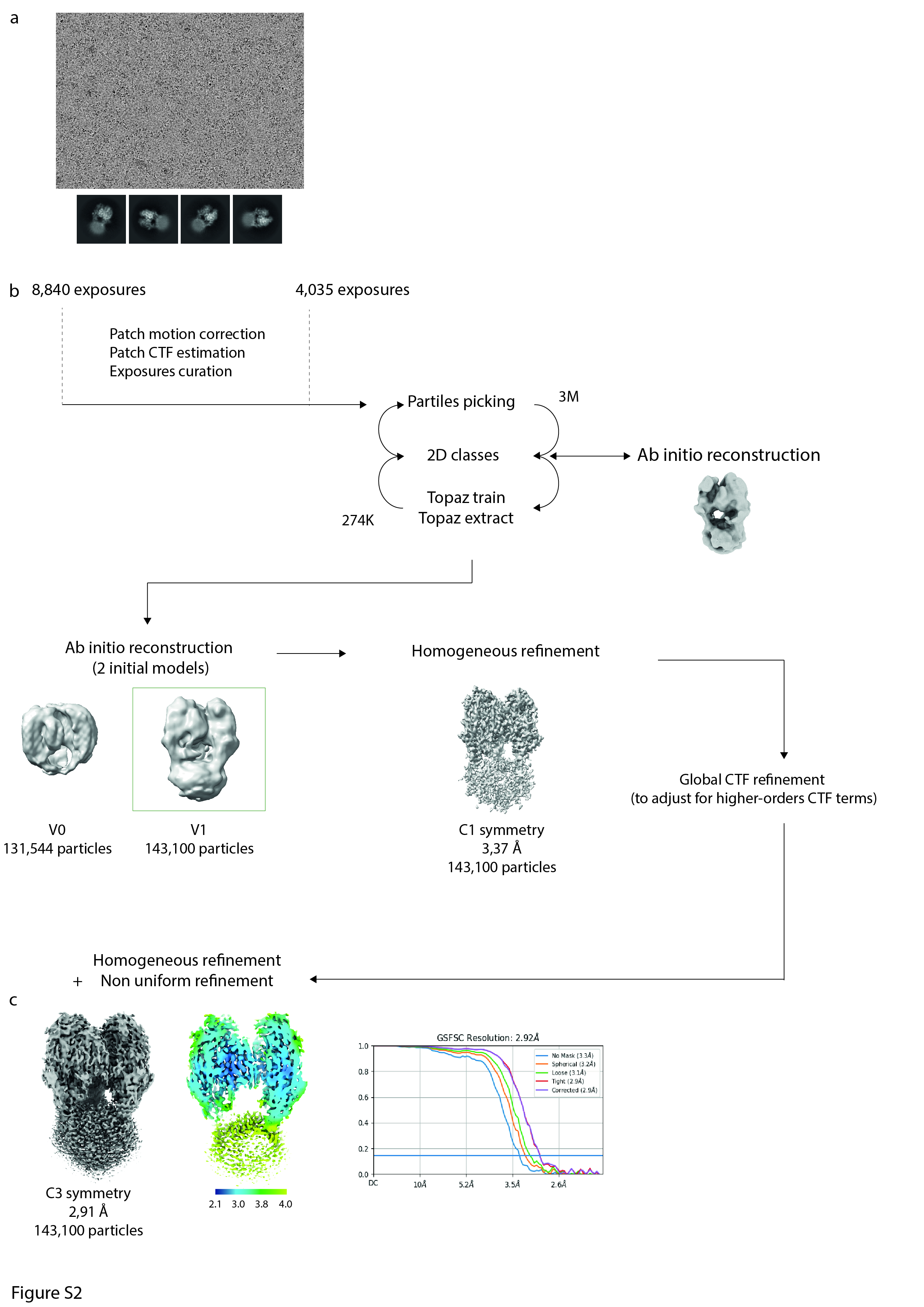

### Supplementary Fig. 3

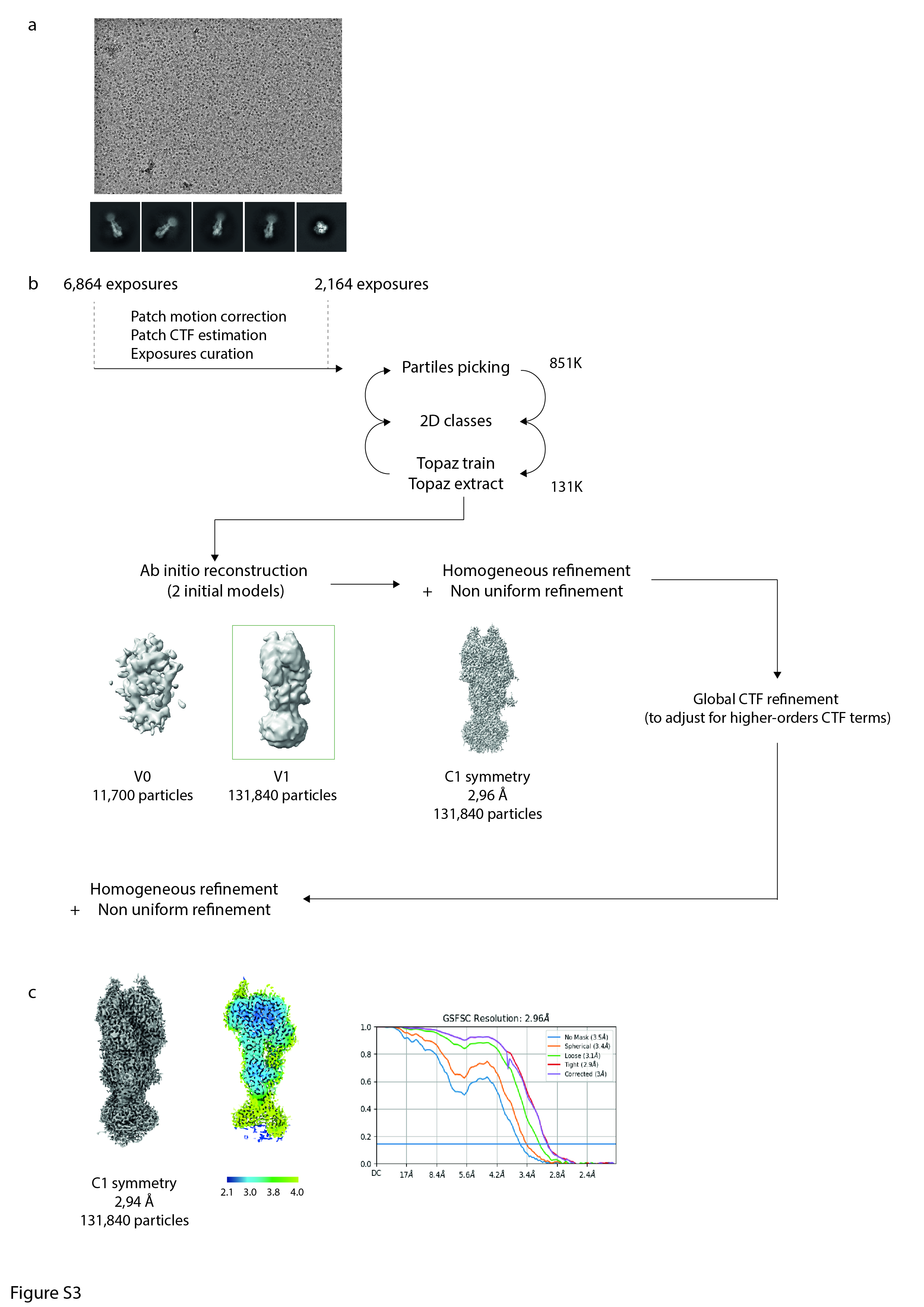

### Supplementary Fig. 4

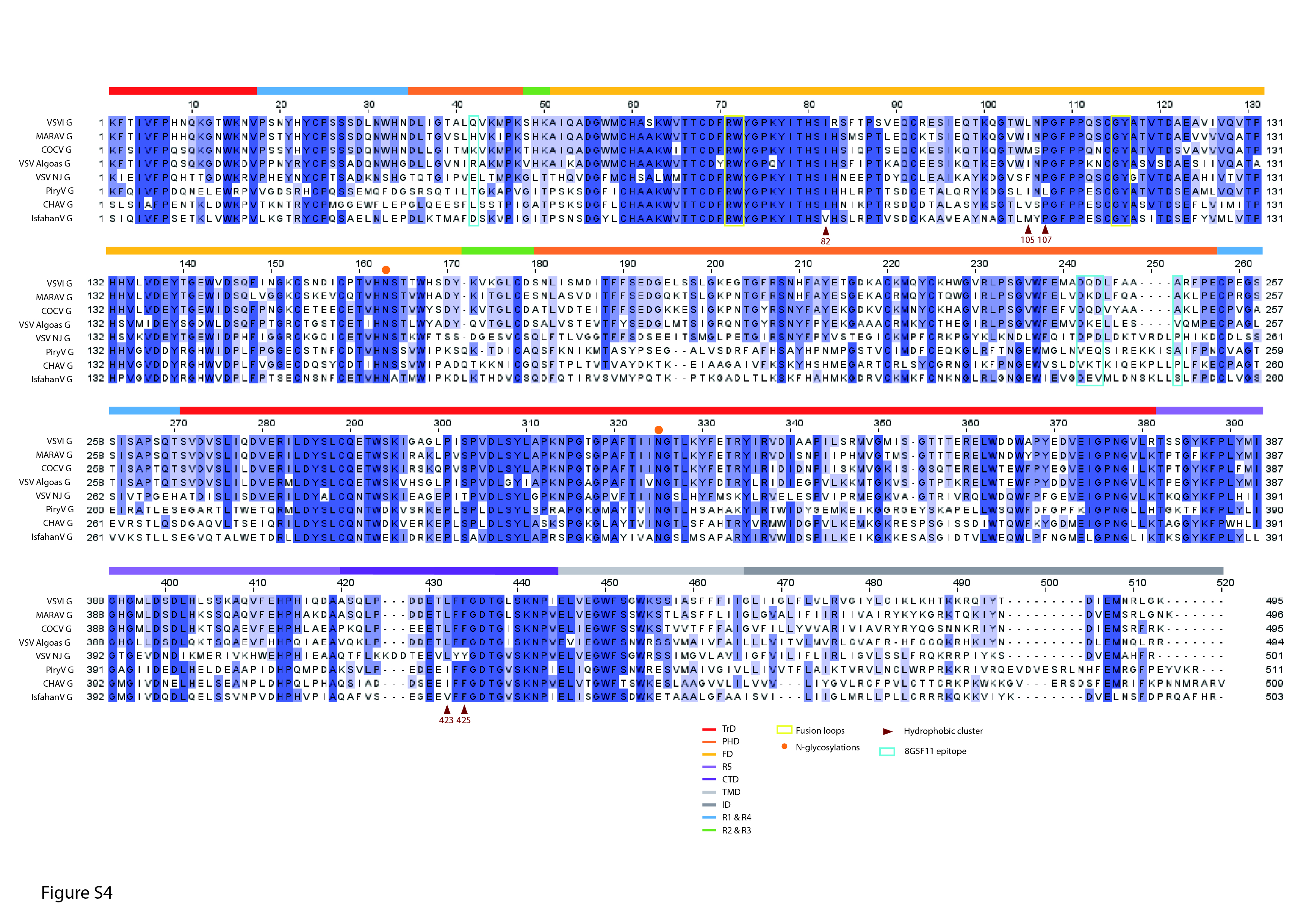

### Supplementary Fig. 5

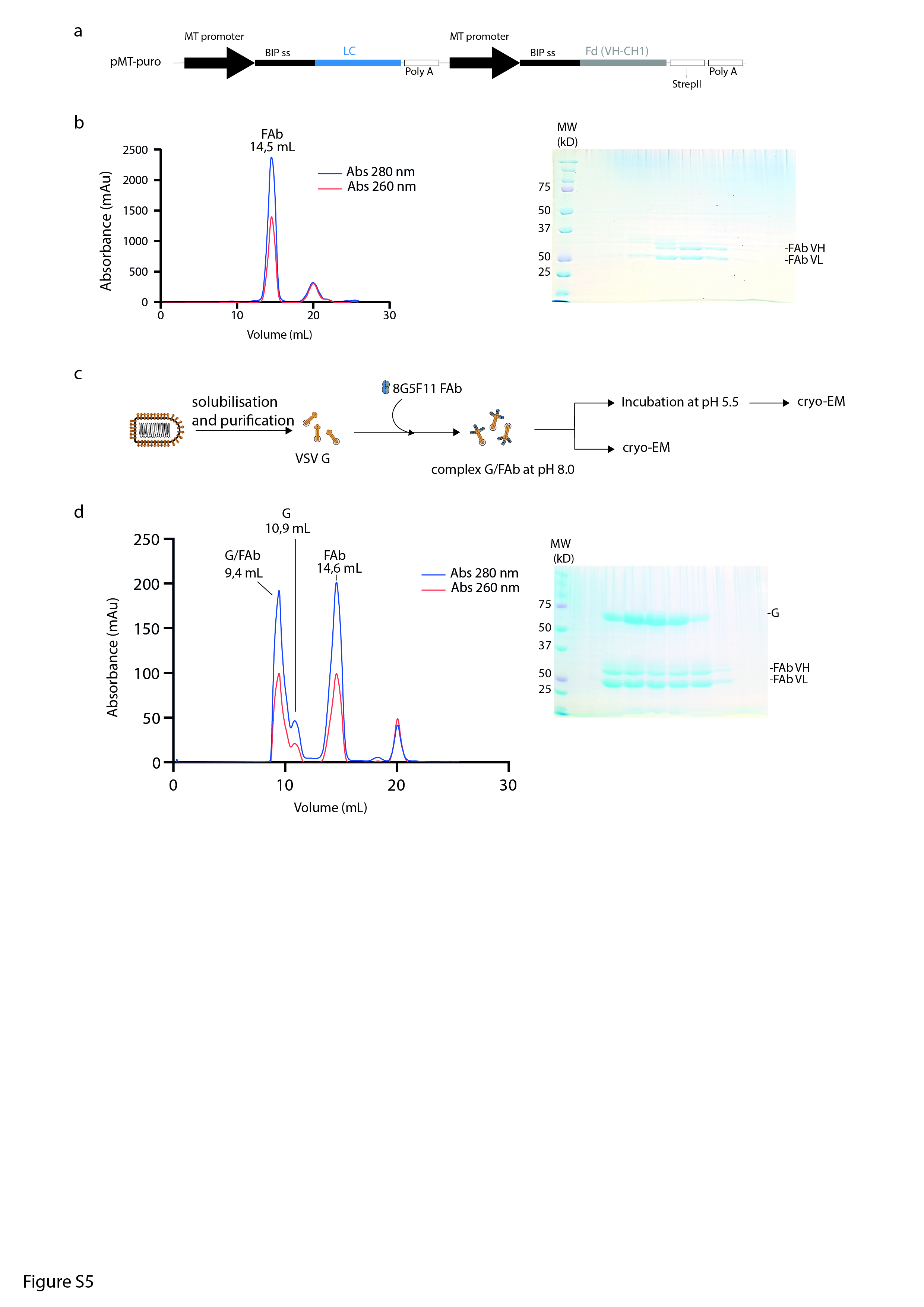

### Supplementary Fig. 6

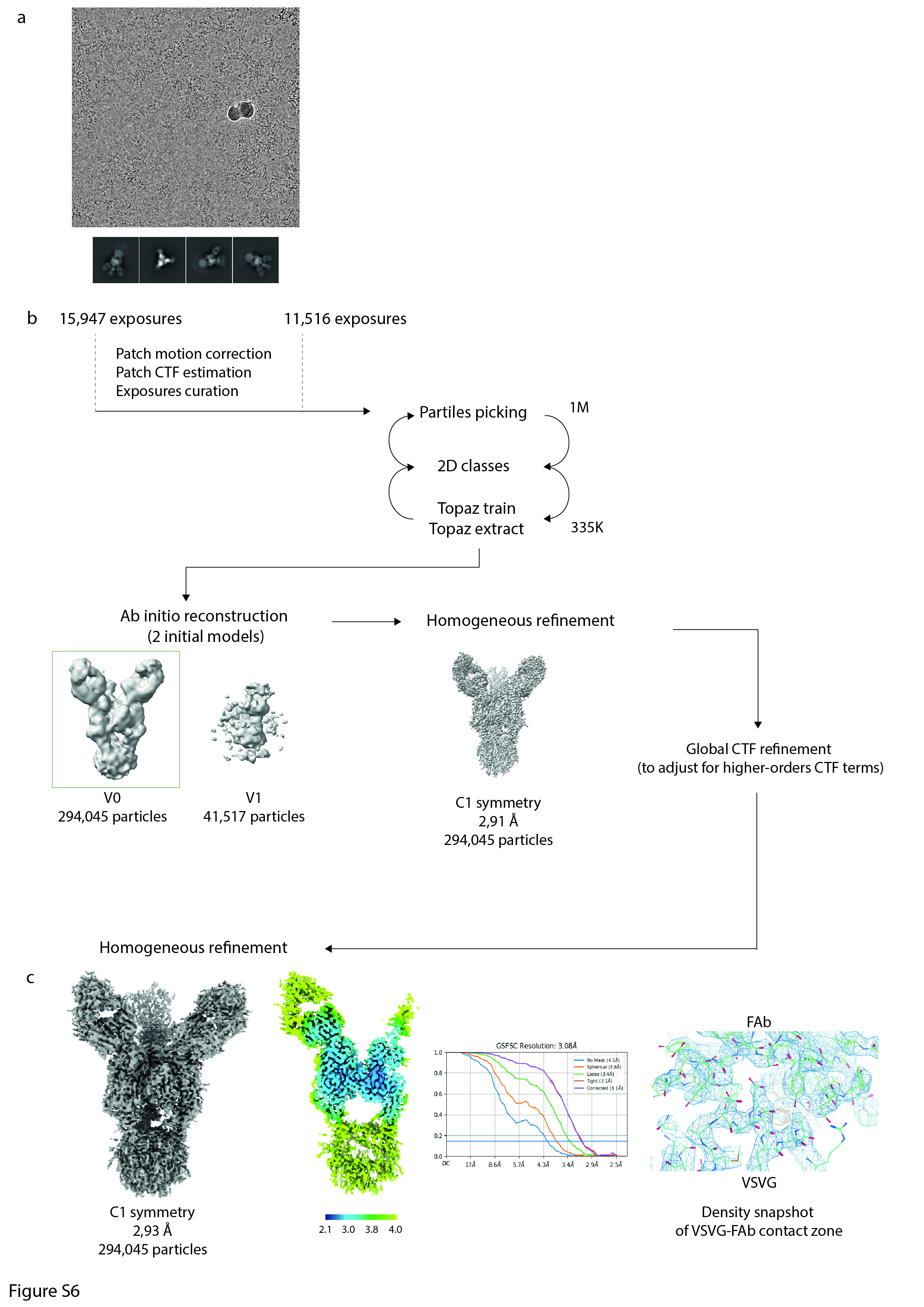

### Supplementary Fig. 7

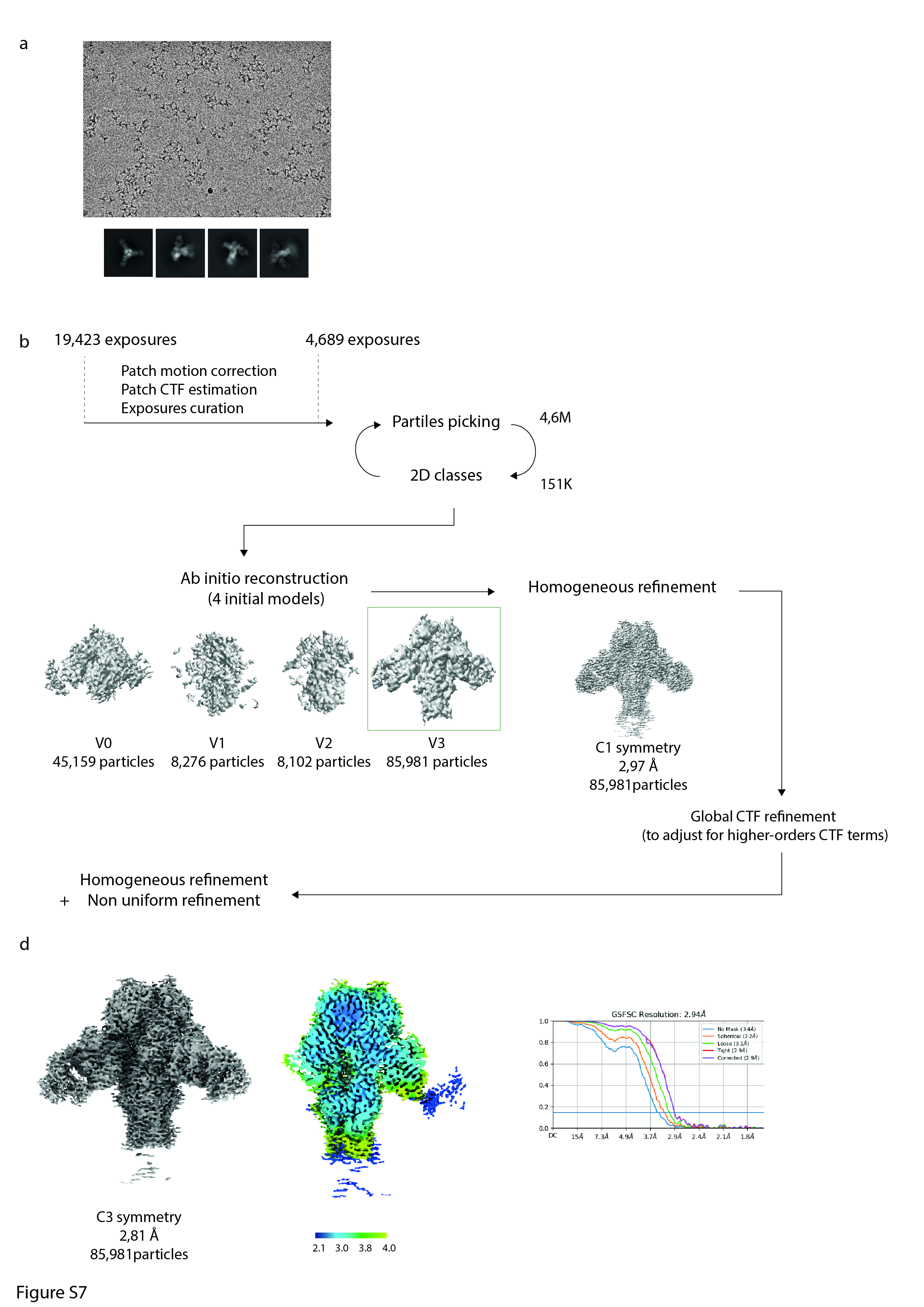

### Supplementary Fig. 8

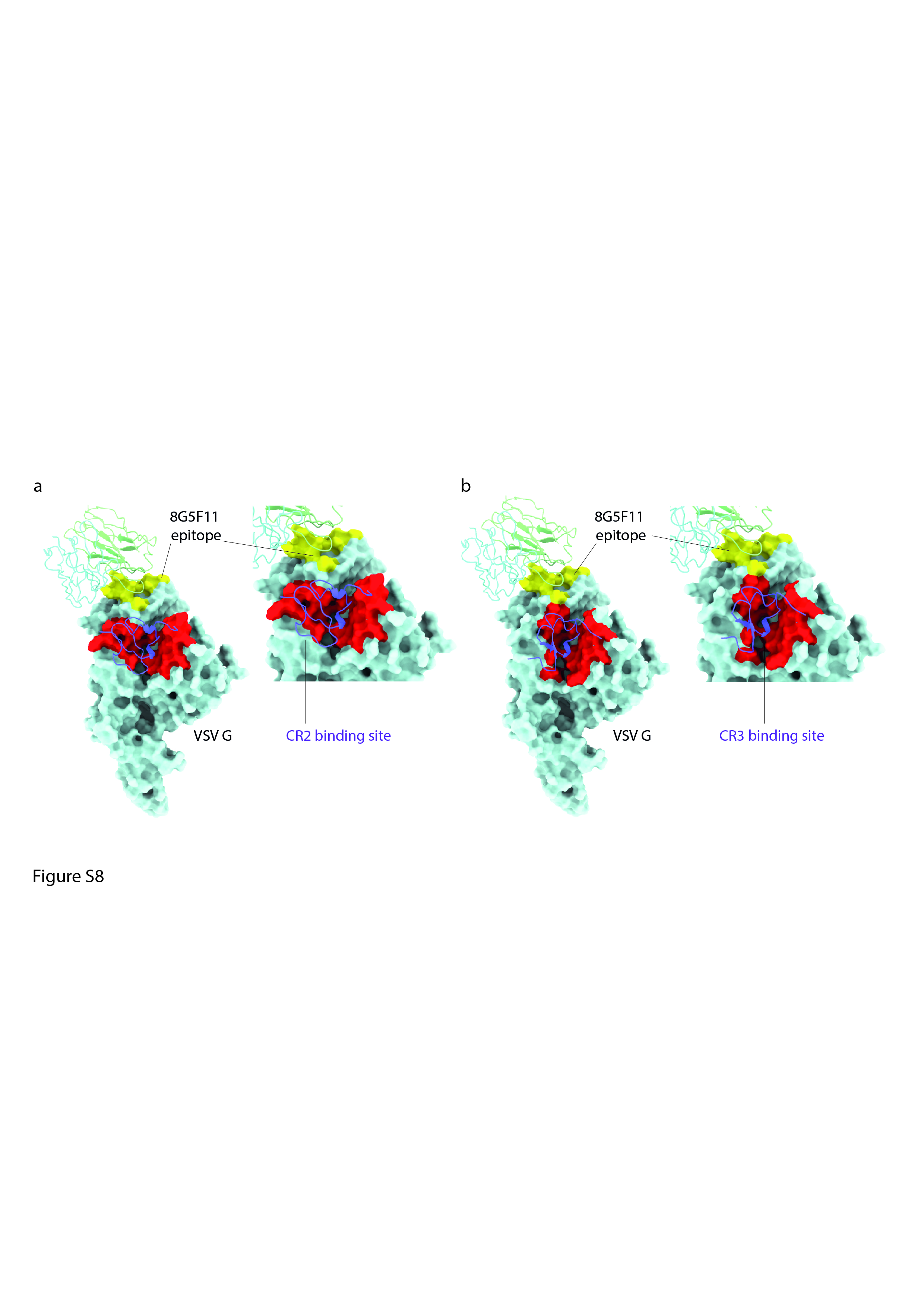
