## Supplementary material for "Structures of vesicular stomatitis virus glycoprotein G alone and in complex with a neutralizing antibody": Table S1

**Table S1. Model and map statistics.**

|  | **VSV Gfl pH 5.0** | **VSV G pH 8.0** | **VSV G – FAb pH 5.0** | **VSV G – FAb pH 8.0** |
| --- | --- | --- | --- | --- |
| **Data collection** | | | | |
| Microscope | Titan Krios | Titan Krios | Titan Krios | GLACIOS |
| Detector | K3 direct electron detector | K3 direct electron detector | K3 direct electron detector | Falcon II |
| Voltage (kV) | 300 | 300 | 300 | 200 |
| Magnification | 105 K | 105 K | 105 K |  |
| Electron exposure (e^-^/Å²) | 40.4 | 40.4 | 49.71 | 38.50 |
| Number of frames | 40 | 40 | 40 | 40 |
| Defocus range (µm) | -0.8 to -2.2 | -0.8 to -2.2 | -0.8 to -2.2 | -0.8 to -2.2 |
| Pixel size (Å/px) | 1.05 | 1.05 | 0.84 | 1.145 |
| **Image processing** | | | | |
| Initial particle images (no.) | 6864 | 8840 | 19693 | 15947 |
| Final particle images (no.) | 4035 | 4890 | 7153 | 6858 |
| Symmetry imposed | C1 | C3 | C3 | C1 |
| Final number of particles | 131.840 | 143.100 | 85.981 | 294.045 |
| Map resolution (Å)  FSC threshold | 2.94  0.143 | 2.91  0.143 | 2.81  0.143 | 2.93  0.143 |

|  | **VSV Gfl pH 5.0** | **VSV G pH 8.0** | **VSV G – FAb pH 5.0** | **VSV G – FAb pH 8.0** |
| --- | --- | --- | --- | --- |
| **Model refinement** | | | | |
| Model to map resolution in Å (FSC 0.5) | 2.98 | 3.02 | 2.93 | 3.28 |
| Map sharpening B-factor (Å) | 88.6 | 128.2 | 102.5 | 81.4 |
| *Model composition*  Non-hydrogen atoms  Protein residues |  | 10092  1278 | 15492  1959 | 15613  1959 |
| *R.m.s deviations*  Bond lengths (Å)  Bond angles (°) | 0.004 (2)  0.664 (4) |  |  |  |
|  |  | 0.004 (4)  0.767 (15) | 0.004 (1)  0.725 (25) | 0.004 (1)  0.865 (20) |
| **Validation** | | | | |
| Molprobity score | 2.15 | 2.49 | 2.51 | 2.89 |
| Clash score | 12.06 | 13.82 | 11.91 | 14.61 |
| Poor rotamers (%) | 2.06 | 2.69 | 4.30 | 6.17 |
| *Ramachadran plot*  Favored (%)  Allowed (%)  Outliers (%) | 95.44  4.48  0.08 | 91.27  7.70  1.02 | 93.25  5.77  0.98 | 87.07  12.00  0.93 |
