## Supplementary material for "Structures of vesicular stomatitis virus glycoprotein G alone and in complex with a neutralizing antibody": Table S2

**Table S2. Average RMSD between aligned Cα (in A) of the overall structure and different regions.**

| **Structural domain** | **Number of residues** | RMSD  VSV G post-fusion / 5i2m | RMSD VSV G pre-fusion / 6tit |
| --- | --- | --- | --- |
| **Whole molecule** | 426 | 0,664 (366 Cα) | 0,634 (369 Cα) |
| **FD** | 120  (res 53 to 173) | 0,549 (108 Cα) | 0,500 (99 Cα) |
| **PHD** | 92  (res 35 to 47 and 180 to 260) | 0,492 (88 Cα) | 0,431 (82 Cα) |
| **TrD** | 127  (res 1 to 18 and 272 to 382) | 0,430 (119 Cα) | 0,403 (121 Cα) |
